# Hippocampal replay anticipates the hexagonal axes that structure human navigation behaviour

**DOI:** 10.1101/2023.02.19.529130

**Authors:** Bo Zhang, Siqi Tu, Jia Liu

## Abstract

Entorhinal grid cells provide a spatial metric through hexagonally periodic firing, whereas hippocampal replay reactivates experience in compressed sequences; whether the two share a directional geometry is unknown. We recorded magnetoencephalography during a direction-resolved navigation task that yielded 1,800 navigation events per participant, allowing both to be estimated event-wise across the full 360°. Reverse and forward replay emerged early in each event, followed by a six-fold entorhinal code. Replay was anchored to the grid axes, with forward and reverse replay encoding opposite directions of each axis and together spanning the six-fold structure. In a continuous-attractor network, replay along the hexagonal axes produced higher gridness than movement-axis or no replay. Behaviourally, an early modulation of movement speed was followed by a two-fold modulation of trajectory deviation whose phase aligned to the hexagonal axes. These results link a single hexagonal geometry across replay, entorhinal coding, and behaviour.

---

The internal map of space in the mammalian brain rests on two phenomena of the hippocampal-entorhinal system. In the hippocampus (HPC), place cells fire at specific locations and are reactivated in temporally compressed sequences, termed replay, in rodents and humans^1,2,3,4,49^. Replay carries distinct functional roles. Forward replay supports prospective, model-based decision-making^5,6^ and reshapes abstract cognitive maps offline^7^, whereas reverse replay re-enacts recently experienced trajectories and propagates value to support reward-based learning^1,8,9^. In the entorhinal cortex (EC), grid cells fire at the vertices of a hexagonal lattice and provide a metric that integrates discrete locations into a continuous map^10^. This metric generalizes beyond physical space to conceptual and even olfactory dimensions^11,12^. Although both phenomena are central to spatial cognition, how the hexagonal grid metric is constructed and stabilised, and whether HPC replay participates in organising it, remains unresolved.

Converging evidence indicates that the HPC shapes the entorhinal grid pattern. The grid pattern is not self-sustaining and requires continuous excitatory drive from the hippocampus. Reducing hippocampal theta input degrades grid periodicity^13^, and inactivating the HPC abolishes grid firing^14^. Hippocampal replay is observed during both rest/sleep and active/awake behaviour^1,3^. The entorhinal grid network participates in these reactivation events. During rest, entorhinal and hippocampal populations replay coherently, with the HPC leading by approximately 11 ms^15^, and superficial entorhinal layers replay independently of the HPC^16^. A developmental dissociation points in the same direction, as preconfigured hippocampal sequences are available early^17^ whereas grid cells mature only after place and head-direction cells^18^. In humans, hippocampal ripples during post-learning rest predict the later emergence of grid-like activity in EC and prefrontal cortex^19^. These findings raise the question of whether replay is organized with respect to the geometry of the grid axes, and whether such coordination is also expressed during active navigation.

Grid orientation is not arbitrary. Grid cells within an individual share a common set of axes, anchored to environmental geometry and preserved across environments^10,20–22^. Entorhinal activity and navigation are both preferentially organized along these axes: conjunctive grid-by-direction cells are tuned to them, and movement along them evokes the strongest population activity^23,24^. Whether replay is likewise organized along these axes, rather than only along the path just travelled, has not been examined. We reasoned that, if replay contributes to the grid metric, it should be anchored to these same axes. Grid axes are undirected, each spanning two antipodal directions, whereas replay is directional: reverse replay retraces a trajectory backwards, so its decoded heading is opposite to that of forward replay^1,2^. We predict that forward and reverse replay should occupy complementary halves of the hexagonal lattice, each covering three of the six grid-aligned directions and the two triplets being antipodal. Each replay type would then be direction-asymmetric on its own, and the six-fold structure should emerge only when the two are combined.

Testing this geometry requires capturing fast replay together with the grid code within the same task, a combination that has remained difficult to achieve with conventional paradigms. Estimating grid geometry requires dense sampling of directions. Rodent electrophysiology resolves replay at the millisecond timescale, but directional sampling is constrained by the animal’s own trajectories. Linear and maze tracks afford only a few headings, whereas in open arenas heading varies continuously along each trajectory and accumulates non-uniformly over hours^1,2^. Human fMRI, which resolves the six-fold grid activity^11,24,25^, lacks the temporal resolution to recover replay sequences. A way forward is offered by the observation that entorhinal grid-like activity is engaged not only during locomotion but also during visual and gaze-based exploration of space^25,26,27,28^. Combined with the millisecond resolution of magnetoencephalography (MEG) and time-lagged decoding of replay^4^, repeated and direction-controlled movements of a visual cursor can in principle evoke entorhinal grid-like activity and hippocampal replay within the same brief and ongoing navigation events, while accumulating enough trial-unique directions to recover their geometry.

We designed an MEG paradigm in which participants navigated between four non-meaningful symbols on a spherical viewing surface using a MEG-compatible mouse system (Fig. 1a–c). The cursor was permanently fixed at the centre of the screen, analogous to the first-person view in a shooter video game, so that the only externally controlled variable was the direction the participant rotated toward. The four symbols were never displayed simultaneously. Each trial began with a fixation period during which participants were already oriented toward the first symbol, and a single click initiated three sequential connections (A→B, B→C, C→D; Fig. 1b). Across five scanning sessions, participants unknowingly sampled 120 uniformly distributed directions for each connection, yielding 1,800 navigation events per participant (5 sessions × 120 directions × 3 connections), allowing replay and grid activity to be estimated event-wise within a single coordinate space.

**Figure 1.**
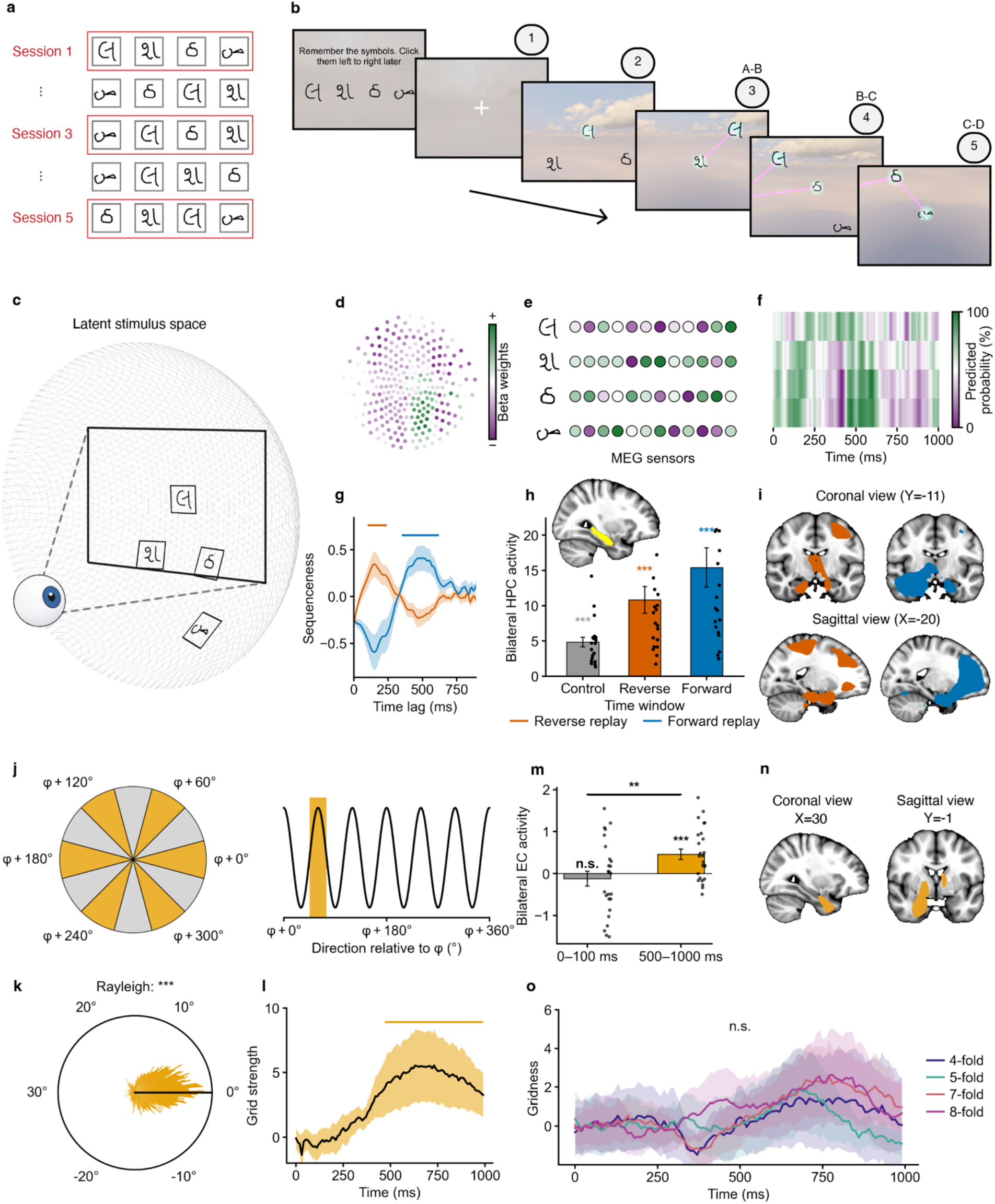
Task Paradigm, hippocampal replay, and entorhinal grid-like coding. (a) Four non-meaningful Omniglot stimuli, used across five sessions; each trial sampled one of 120 predefined movement directions. (b) Example trial: instruction cue → fixation (1) → the first symbol presentation (2)→ three sequential pair connections (A→B, B→C, C→D; 3-5). (c) Schematic of the latent stimulus space. Stimuli embedded in a two-dimensional latent space, only a portion was visible at any time and the four stimuli were never shown simultaneously. Participants navigated with a screen-centred cursor under a self-centred reference frame. (d-e) Schematic of sensor-level beta-weight topographies estimated from a one-vs-rest logistic-regression decoder. (f) Schematic of state-to-state reactivation map. (g) Reverse and forward replay reached significance around 150 ms and 500 ms (reverse: 100–240 ms; forward: 360–620 ms; cluster-based permutation p < 0.05, one-tailed). (h) Bilateral HPC activity in control (0–100 ms), reverse and forward windows; both replay windows exceeded control (***p < 0.001; mean ± s.e.m.). (i) Reverse and forward replay localised to bilateral HPC, extending into frontal lobe (cluster-based permutation p < 0.05, two-tailed). (j) Six-fold (60°-periodic) sinusoidal modulator. (k) Grid-like code in EC: within-subject voxel-coherence R = 0.38 ± 0.05, Rayleigh test p<0.001. (l-m) The six-fold grid signal was significant in the late window (500–1000 ms), exceeding the early window (t(24) = 2.82, **p = 0.009). (n) source-level clusters in bilateral EC at 500–1000 ms. (o) Control: adjacent folds (4-, 5-, 7-, 8-fold) showed no significant grid signal (all p > 0.05).

## Results

### A direction-resolved latent-navigation task elicits hippocampal replay and entorhinal grid-like activity

All participants completed the connection task with high accuracy (Fig. S1a; mean accuracy = 99.22% ± 0.10% s.e.m.), and motion directions tightly aligned with the veridical direction (Fig. S1c & Fig. S3-4; circular mean direction error = −1.73°, circular s.d. = 3.7°; Rayleigh test, p < 1 × 10⁻¹²). Each connection lasted approximately 1 s (Fig. S1b; Pair-AB = 987 ± 59 ms; Pair-BC = 957 ± 43 ms; Pair-CD = 968 ± 44 ms; F(2, 48) = 0.46, p = 0.634), motivating the 1-s analysis window adopted in subsequent analyses.

To detect hippocampal replay activity, a one-vs-rest logistic-regression decoder was trained at each 10-ms timepoint for each stimulus (Fig. 1d & e). Stimulus identity was reliably decodable: averaging across the four classifier–class diagonals, decoding peaked at ∼300 ms post-stimulus (t(24) = 6.32, p = 1.5 × 10⁻⁶), and for each stimulus the correct class significantly exceeded the mean of the other three classes across a sustained window (Fig. S2; Bonferroni-corrected across timepoints, p < 0.05). The decoder trained at the peak-accuracy timepoint was then applied to MEG activity during the connections, yielding stimulus-specific reactivation time-courses (Fig. 1f).

Temporally delayed linear modelling (TDLM) revealed significant reverse and forward replay with distinct temporal profiles (Fig. 1g; initial threshold p < 0.05, cluster-based permutation p < 0.05, one-tailed). Reverse replay emerged early, peaking around 150 ms (cluster: 100–240 ms). Forward replay emerged later, peaking around 500 ms (cluster: 360–620 ms).

To validate replay activity within the early and late time window, the bilateral HPC activity for both replay types was compared with that in a control window (0–100 ms). All three windows showed significant activity in the HPC (Fig. 1h; control [0–100 ms]: t(24) = 7.22; reverse [100– 240 ms]: t(24) = 5.72; forward [360–620 ms]: t(24) = 5.51; all p < 0.0001), and the reverse and forward windows showed significantly stronger activity than the control window (reverse vs control: t(24) = 3.97, p = 8 × 10⁻⁴; forward vs control: t(24) = 4.38, p = 3 × 10⁻⁴). Whole-brain source localization confirmed the replay activity in the bilateral HPC (Fig. 1i; initial threshold p < 0.001, cluster-based permutation p < 0.05, two-tailed), which extended into the medial and lateral frontal lobe and motor cortex.

We next tested whether co-occurring entorhinal grid-like activity could be detected during these same connections. Following the established approach with a five-fold cross-validation procedure (see Methods for details), we aligned a six-fold sinusoidal modulator based on each participant’s φ (Fig. 1j) and tested for a population-level six-fold modulation of EC activity as a function of direction.

Within each participant, grid orientation φ was significantly clustered across EC voxels (Fig. S5; per-participant Rayleigh test), with a mean within-subject R = 0.38 ± 0.05 s.e.m. After calibrating each participant’s φ to its own mean orientation, the pooled distribution remained significantly non-uniform within the 60° wrapping window (Fig. 1k; Rayleigh test, p < 0.001).

Within the bilateral EC, a significant cluster of six-fold grid-like activity emerged at 500– 1000 ms after connection onset (Fig. 1l; initial threshold p < 0.05; cluster-based permutation p < 0.05, one-tailed), and showed significantly higher activity strength above zero (t(24) = 3.67, p = 6 × 10⁻⁴), whereas no significant activity was observed in the early control window (0–100 ms; t(24) = −0.68, p = 0.75). This late-window activity was also significantly stronger than that in the early window (Fig. 1m; t(24) = 2.82, p = 0.009). Moreover, source-level analysis revealed clusters in bilateral EC at 500–1000 ms (Fig. 1n; initial threshold p < 0.001, cluster-based permutation p < 0.05, two-tailed). As a control, none of the adjacent control folds (4-, 5-, 7-, 8-fold) showed a significant grid signal (Fig. 1o; p > 0.05).

### Hippocampal replay activity aligns with the later entorhinal grid orientation

The observation that replay activity (reverse: 100–240 ms; forward: 360–620 ms; Fig. 1g) preceded EC six-fold activity (500–1000 ms; Fig. 1l) raised the question of whether hippocampal replay events are spatially anchored to the entorhinal grid metric that emerges later. To address this, we tested whether reverse and forward replay were aligned to the grid axes in the directional domain. In doing so, replay activity was estimated for each of 120 directions (Fig. 2a, II), giving each connection a direction-by-direction replay profile. The direction label of replay activity was then realigned to the participant’s grid orientation φ. Every direction was shifted by −φ and grid-aligned directions (multiples of 60° from φ) and grid-misaligned directions (offsets of 30°) became comparable across participants (Fig. 2a, I). Next, replay activity was contrasted between grid-aligned and grid-misaligned directions (Fig. 2a, III). Given that the EC grid periodicity emerges after replay (Fig. 1g & l), the window in which grid orientation is estimated need not coincide with that in which replay is measured; we therefore examined all combinations of the two: each replay time-point was realigned using the grid orientation extracted at every grid-orientation time-point (Fig. 2a, IV). Throughout, grid orientation φ was estimated under five-fold cross-validation (fit on four folds, evaluated on the held-out fold) and replay was measured on held-out trials, so that the φ used to define grid-aligned versus grid-misaligned directions was obtained from cross-validated estimates rather than from the same data that produced the replay measure.

**Figure 2.**
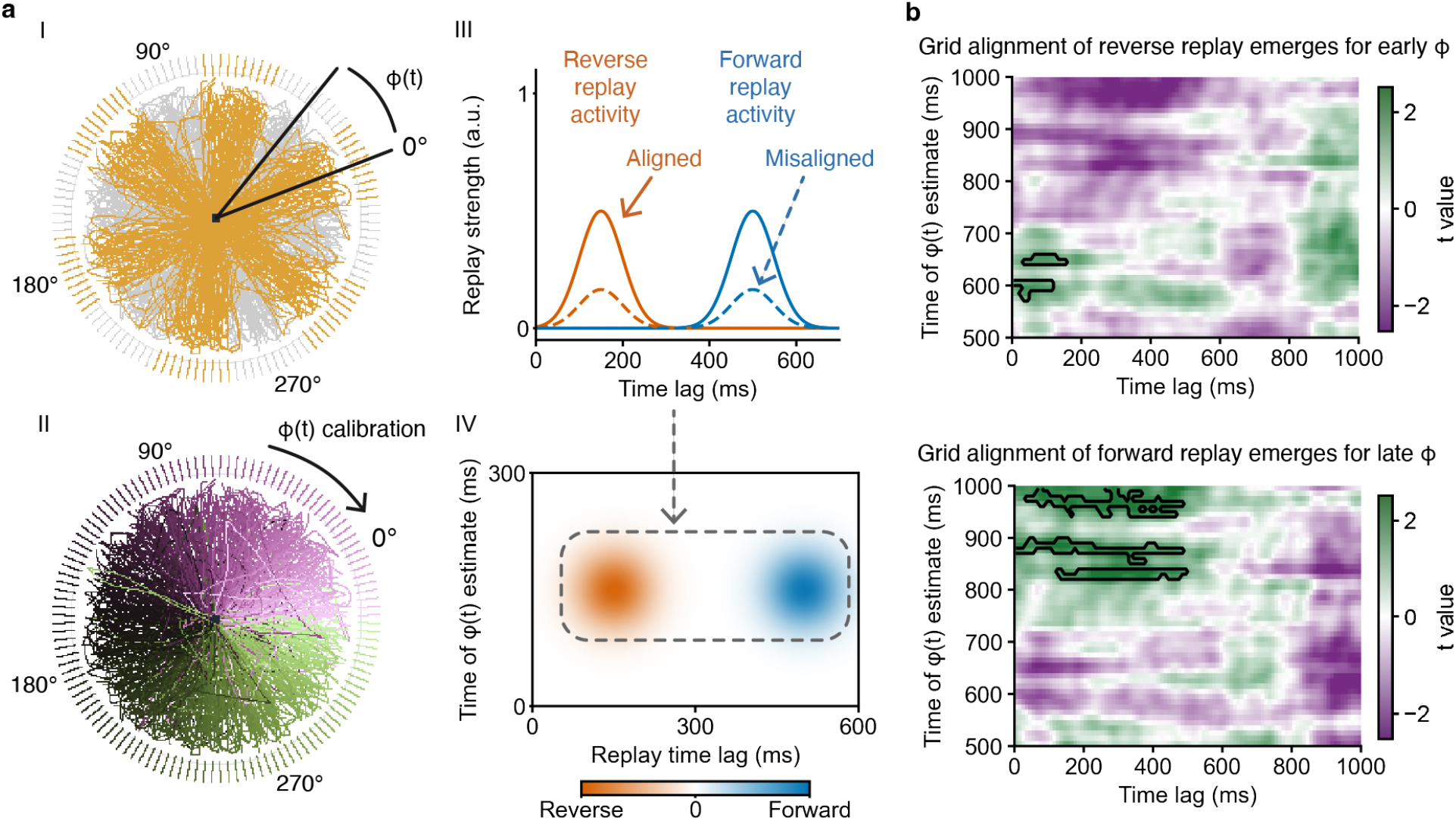
Replay is aligned with entorhinal grid orientation. (a) Analysis pipeline for the replay– grid alignment test. (I) For a representative participant, the entorhinal grid orientation φ (estimated at a given grid-orientation time t) defines the three grid axes, and labels each movement direction as grid-aligned (orange; within ±15° of a multiple of 60° from φ(t)) and grid-misaligned (grey). (II) Replay activity is estimated separately for each of the 120 movement directions (color denotes direction); the per-direction sequence is then rotated by −φ(t) so that the replay activity in grid-aligned and grid-misaligned directions are comparable across participants. (III) Aligned−misaligned contrast for each type of replay activity. (IV) Schematic 2D align − misalign contrast map over replay time lag (x axis) and φ-estimation time (y axis); each row shows the align − misalign replay contrast across replay time, calibrated against the grid orientation estimated at one φ-estimation time point. (b) Cluster-based statistical maps showing where replay activity differed between the aligned and misaligned conditions, as a function of replay analysis time (x-axis) and grid-orientation calibration time (y-axis). Top, reverse replay; bottom, forward replay. Colour indicates t value. Black contours mark significant clusters (initial threshold p < 0.05, cluster-based permutation p < 0.05, one-tailed). Reverse replay showed a significant cluster at early replay times and earlier grid-orientation calibration windows; forward replay showed a significant cluster at later replay times.

The resulting align − misalign difference map identified significant replay-time - grid-orientation-time pair clusters for both replay conditions (Fig. 2b; initial threshold p < 0.05, cluster-based permutation p < 0.05, one-tailed). Reverse replay showed significantly higher strength in the aligned than in the misaligned condition at around 150 ms (replay time) when realigned by the 570–650 ms grid-orientation window, and forward replay showed the same align > misalign difference at a later time window than reverse replay, extending to 490 ms when calibrated by the 820–990 ms grid-orientation time window. These results suggested that reverse replay (peaking ∼150 ms) was followed first by an earlier (570–650 ms) grid-orientation alignment, while forward replay (peaking ∼500 ms) was followed by a later (820–990 ms) grid-orientation alignment, mirroring the order of the replay events in Fig. 1g, and suggesting that grid-axis orientation time-locked to the preceding replay events.

### Forward and reverse replay form complementary half-lattices on shared grid axes

We next asked whether replay activity was specifically organized by the hexagonal grid geometry. A six-fold pattern would be expected if replay directions align with the grid axes, with grid-aligned bins showing higher activity. Strikingly, replay activity binned into 12 directions relative to φ deviated from this full-six-axis prediction (Fig. 3a). For reverse replay, the three high-amplitude grid-aligned bins were found at 0°, 60° and 120°, exceeding the grid-misaligned baseline (t(24) = 4.63, p = 1.1 × 10⁻⁴, Cohen’s d = 0.93), whereas the other three grid-aligned bins (180°, 240°, 300°) with opposite-direction to 0°, 60° and 120° did not differ from the grid-misaligned baseline (t(24) = 0.32, p = 0.75, d = 0.06). Forward replay significantly coded three grid-aligned bins at 180°, 240° and 300°, exceeding the grid-misaligned baseline (t(24) = 4.50, p = 1.5 × 10⁻⁴, d = 0.90), whereas the three opposite grid aligned directions (0°, 60°, 120°) did not differ from the grid-misaligned baseline (t(24) = 0.96, p = 0.35, d = 0.19). Forward and reverse replay therefore showed mutually exclusive direction selectivity. Each was tuned to three of the six grid-aligned directions, and the two preferred triplets pointed in opposite directions (180° apart) along the same three hexagonal axes.

**Figure 3.**
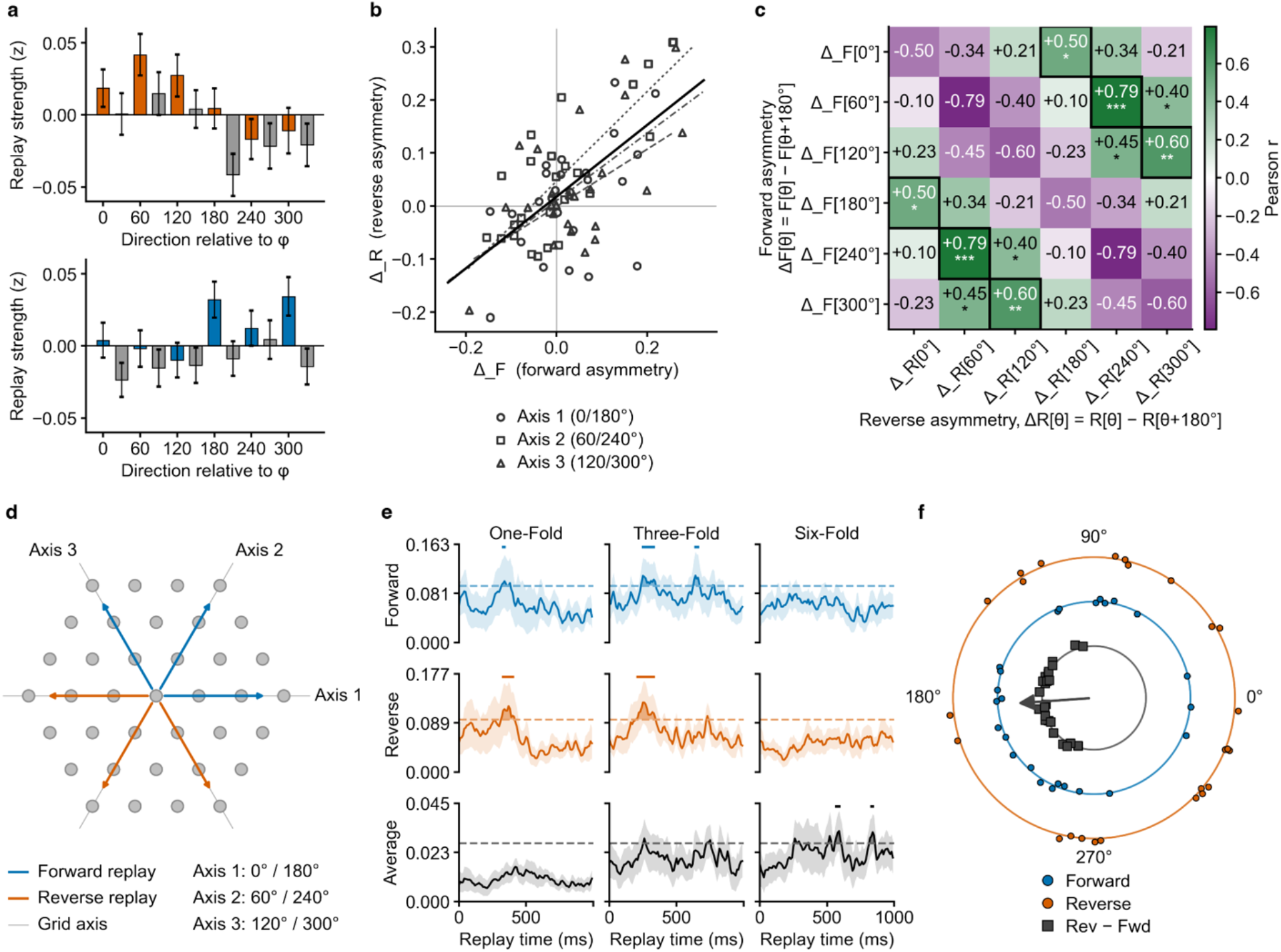
Forward and reverse replay traverse opposite directions on grid axes. (a) Selective direction tuning of replay shown separately for reverse (top) and forward (bottom) replay. Color bars indicate align directions; grey bars indicate misalign directions. Replay activity in the selective directions significantly exceeded the misaligned baseline in both conditions (forward: t(24) = 4.50, p = 1.5 × 10⁻⁴; reverse: t(24) = 4.63, p = 1.1 × 10⁻⁴, two-tailed; Cohen’s d = 0.90 and 0.93). (b) Forward and reverse replay co-vary on grid axes. Left, across-participant correlation: forward replay at each direction was correlated with reverse replay at the opposite direction (forward directional asymmetries: Δ_F; reverse directional asymmetries: Δ_B; axis 1 [0°/180°]: r = 0.50, p = 0.011; axis 2 [60°/240°]: r = 0.79, p = 2.2 × 10⁻⁶; axis 3 [120°/300°]: r = 0.60, p = 0.001). (c), full 6 × 6 across-participant correlation matrix between forward and reverse directional asymmetries across the six grid-aligned directions; cells show Pearson r (*p < 0.05, **p < 0.01, ***p < 0.001). Bold-bordered cells mark the six ’shared-axis, opposite-direction’ pairs (the shift-3 diagonal, where forward replay at θ pairs with reverse replay at θ + 180°), which showed significantly higher correlation than the remaining 30 cells (paired t-test, t(24) = 3.47, p = 0.002, two-tailed; Cohen’s d = 0.69). (d) Schematic summarising that forward and reverse replay span all six grid-aligned directions, with each replay type traversing only one of the two opposite directions on each axis. (e) FFT power of grid-orientation–realigned replay activity as a function of replay time, shown for forward (top), reverse (middle) and the forward–reverse average (bottom), at folds 0–6. Forward and reverse replay each showed significant one- and three-fold power but no six-fold power, whereas the average showed significant six-fold power within the EC grid-coding window (initial threshold p < 0.05, cluster-based permutation p < 0.05, one-tailed). (f) Per-participant FFT phase (three-fold) of forward and reverse replay in raw motion-direction space. Gray dots indicate per-subject phase difference (reverse − forward). Arrow shows the phase-difference mean resultant (R = 0.76, Rayleigh test, p = 4.2 × 10⁻⁸, circular mean = −176°, expected = 180°).

To test directly whether the three preferred directions of forward replay and those of reverse replay mirror each other on the same grid axes, we computed an individual-participant axis-pair directional asymmetry, defined as the difference in replay activity between the two opposite directions of that axis for each grid axis (i.e., Δ_F[θ] = Forward [θ] − Forward [θ + 180°] and Δ_R[θ] analogously). We therefore predicted that Δ_F and Δ_R would be positively correlated across participants within each grid axis.

Axis-by-axis Pearson correlations across participants revealed significant positive correlations on all three grid axes (Fig. 3b; axis 1 [0°/180°]: r = 0.50, p = 0.011; axis 2 [60°/240°]: r = 0.79, p = 2.2 × 10⁻⁶; axis 3 [120°/300°]: r = 0.60, p = 0.001).

To further validate the axis-by-axis prediction, we extended the analysis to the full 6 × 6 across-participant correlation matrix between forward (rows) and reverse (columns) axis-pair directional differences across the six grid-aligned directions (Fig. 3c). The six “shared-axis, opposite-direction” cells (the shift-3 diagonal, where forward replay at θ pairs with reverse replay at θ + 180°) showed significantly higher correlation strength than the remaining 30 cells (paired t-test, t(24) = 3.47, p = 0.002, two-tailed; Cohen’s d = 0.69).

In sum, the two types of replay activity span all six grid-aligned directions and recover the underlying hexagonal axis structure observed in EC (Fig. 3d). Each replay type individually, however, traverses only one of the two opposing directions on each axis.

We further questioned whether a six-fold periodicity could be detected in replay activity. We predicted that (i) this analysis would reveal significant one- and three-fold power but not six-fold power for forward and reverse replay considered separately (since each replay type was associated with only three of the six grid-aligned directions), and (ii) averaging forward and reverse replay would instead yield six-fold power, with the one- and three-fold components cancelling out. Grid-orientation–realigned forward and reverse replay activity were analysed independently with an FFT, and spectral power at folds 0–6 was tested at each time point.

Both forward and reverse replay reached significance at one-fold and three-fold (Fig. 3e, top and middle; initial threshold p < 0.05, cluster-based permutation p < 0.05, one-tailed). The main clusters fell in the early replay period (forward three-fold, 240–340 ms; forward one-fold, 320–350 ms; reverse three-fold, 200–340 ms; reverse one-fold, 320–410 ms). Forward replay showed an additional later three-fold cluster at 630–670 ms. Neither replay direction showed a significant six-fold cluster individually (p > 0.05), consistent with the direction-asymmetric structure of each replay direction (Fig. 3a).

The averaged activity reached significance primarily at six-fold after 500 ms (Fig. 3e; initial threshold p < 0.05, cluster-based permutation p < 0.05, one-tailed), with clusters at ∼510– 600 ms and ∼820–850 ms, both within the EC six-fold grid-coding window (Fig. 1l; 500–1000 ms). In contrast, one- and three-fold power did not reach significance in the averaged activity (p > 0.05). Overall, the early one-/three-fold replay clusters versus the late six-fold clusters mirror the temporal sequence from hippocampal replay (Fig. 1g) to entorhinal grid coding (Fig. 1l-n).

The anti-phase robustness of forward and reverse replay was further validated at the level of individual-participant phase. Because the direction tuning of replay is anchored to a participant’s own grid orientation φ, which is idiosyncratic, the absolute phase measured in unrealigned motion-direction space carries no consistent across-participant reference and is therefore not expected to be clustered. The within-participant reverse − forward difference, by contrast, cancels φ and should cluster around π if the two replay types are selectively associated with axis-opposite directions.

We extracted and compared the activity phase from individual-participant between both reverse and forward replay. Consistent with our prediction, the absolute three-fold phase was not significantly clustered across participants for either type of replay activity (Fig. 3f; forward: R = 0.27; Rayleigh test, p = 0.15; reverse: R = 0.21; p = 0.33). The reverse − forward phase difference, however, was tightly clustered near π (Fig. 3f; R = 0.76, Rayleigh p = 4.2 × 10⁻⁸; circular mean = −176°). The same anti-phase relation was observed at fold-1 (R = 0.66; circular mean = 167.5°; Rayleigh p = 6.6 × 10⁻⁶).

### Empirically constrained grid-axis replay stabilizes entorhinal grid formation

The empirical findings established that hippocampal replay events are anchored to entorhinal grid axes and that forward and reverse replay share the same hexagonal axes in opposite directions. To test whether this empirically observed replay pattern supports stable grid emergence, we ran a continuous-attractor network (CAN) simulation.

We adopted a CAN implementation that generates grid-cell firing through path-integration input^29^. The network comprised 101 × 101 grid-cell units, with spatial frequency ω determined by the target grid scale σ. To simulate the time compression of empirical replay, the replay-input amplitude was scaled relative to the motion-direction input amplitude (Supplementary Table S1).

Three replay conditions (no-replay, motion-axis replay, grid-axis replay) were tested at 11 grid scales (σ = 20–70 px in 5-px steps) and 12 directions ranging from 0° to 165° in 15° steps; the other half of the directional space, 180°–345°, is symmetric and produces identical CAN dynamics, thus only the upper half was simulated.

In the no-replay condition, the CAN was driven by motion direction input only. In the motion-axis replay condition, replay events were simulated along the motion direction (forward) and its 180° opposite (reverse). In the grid-axis replay condition, forward replay was simulated simultaneously along the three nearest hexagonal axes around the motion direction, while reverse replay was simulated along the 180° opposite hexagonal axes, as observed in the MEG data.

Grid-axis replay produced significant grid-like coding at all eleven grid scales (Fig. 4a, right panel and Fig. 4b; 132/132 simulations significant; mean gridness = 1.32). Motion-axis replay produced significant gridness in 108/132 simulations (81.8%; mean gridness = 1.17; initial threshold p < 0.001, FWE-corrected p < 0.05), and no-replay in 72/132 (54.5%; mean gridness = 0.87); non-significant simulations are shown in greyscale (Fig. 4a). Under no-replay, failures were concentrated at larger spatial scales, with the proportion of significant simulations dropping from 86.1% at small scales (20–45 px) to 16.7% at large scales (50–70 px). Accordingly, final gridness was significantly higher at small than at large scales (1.14 vs. 0.55; paired t-test across the 12 motion directions, t(11) = 7.70, p < 0.001, Cohen’s d = 2.22). Motion-axis replay showed a similar limitation, failing to produce stable hexagonal patterns at large grid scales (σ = 70 px: 0/12 simulations significant; σ = 45 px: 6/12; σ = 40 px: 8/12). With gridness averaged across the 11 scales, motion-axis replay was significantly stronger than no-replay (paired t(11) = 6.56, p = 4.1 × 10⁻⁵, Cohen’s d = 1.90), and grid-axis replay was significantly stronger than motion-axis replay (t(11) = 5.58, p = 1.7 × 10⁻⁴, d = 1.61).

**Figure 4.**
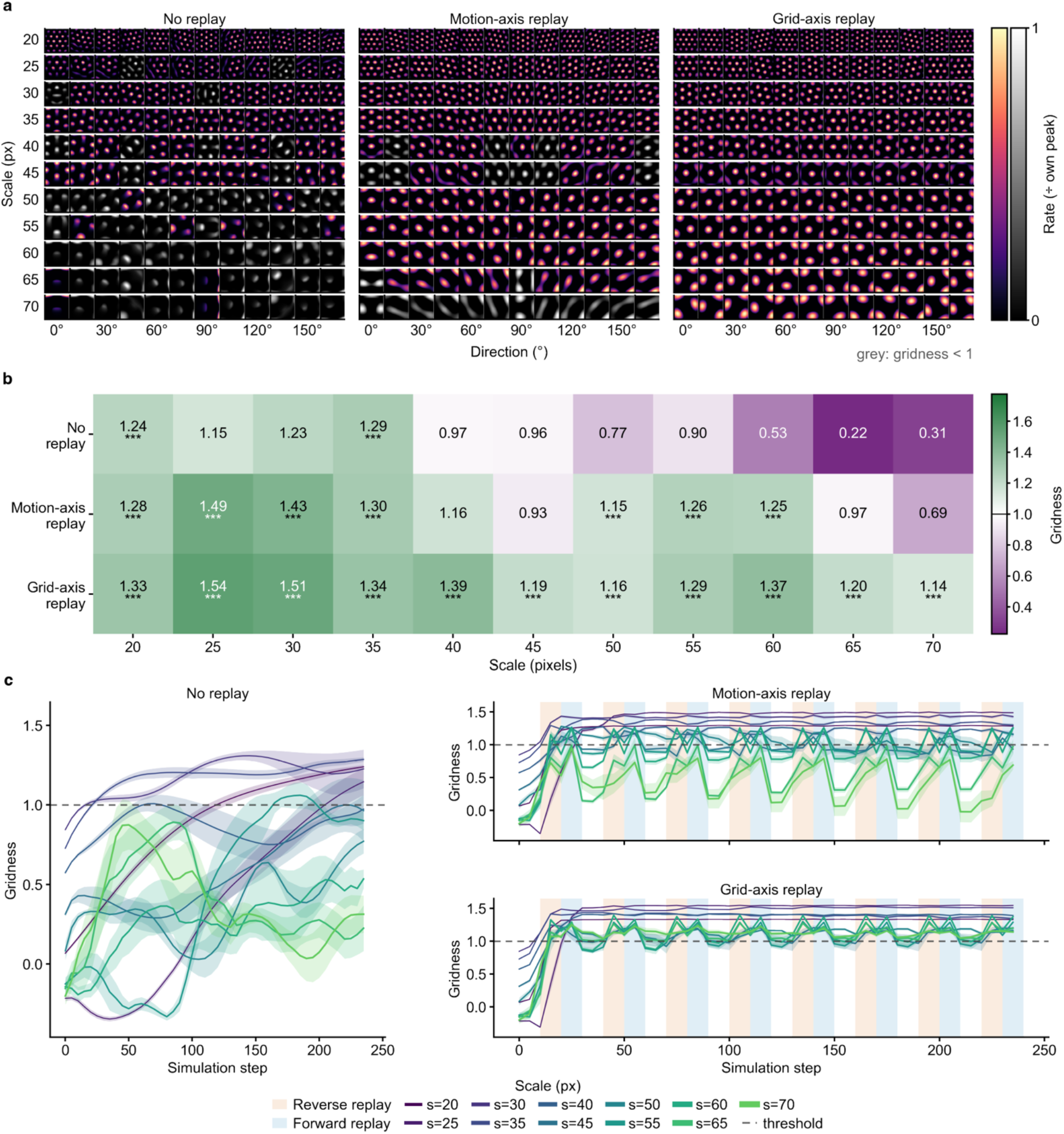
Grid-axis replay supports robust grid formation and accelerates grid emergence in a continuous-attractor network. (a) Grid-like patterns produced by the CAN simulation for every (scale × direction × replay condition) combination. Eleven grid scales (20–70 px in 5-px steps) × twelve directions (0°–165° in 15° steps) for each condition: no replay (left), motion-axis replay (middle; replay along the current motion axis) and grid-axis replay (right; replay along the three hexagonal grid axes). Non-significant gridness was shown in greyscale. (b) Mean gridness across the twelve directions for each combination of replay condition and grid scale (one-sample t-test of mean gridness against threshold = 1.0; Bonferroni-corrected across 33 tests; ***p < 0.001, **p < 0.01, *p < 0.05). Grid-axis replay maintained significant gridness across all eleven scales. (c) Temporal evolution of gridness across simulation steps for each replay condition (mean ± s.e.m. across the 12 directions). Orange and blue bands indicate reverse-replay and forward-replay modulation intervals. Grid-axis replay drives faster and more stable convergence than motion-axis replay or no replay.

Next, we examined how gridness evolved over simulation steps in each of the three conditions (Fig. 4c). Under motion-axis replay, gridness rose during the reverse-replay phase of each cycle and partially returned toward baseline after the forward-replay phase, producing periodic modulation. Under grid-axis replay, gridness rose to higher peak levels (1.38 vs. 1.31; paired t(11) = 6.09, p < 0.001, Cohen’s d = 1.76), with a significantly smaller within-cycle return than under motion-axis replay (modulation amplitude 0.18 ± 0.02 vs. 0.35 ± 0.03; paired t(11) = 15.4, p < 10⁻⁸, Cohen’s d = 4.44), and stable hexagonal patterns were obtained at every grid scale (12/12 directions yielded significant gridness at every σ; initial threshold p < 0.001, FWE-corrected p < 0.05). By contrast, under no-replay condition, gridness rose slowly and aperiodically, and 54.5% of simulations declined after reaching their peak (mean peak-to-trough drop = 0.36 ± 0.04), resulting in a failure to stabilize the emerging grid patterns.

Overall, both replay conditions improved grid formation over the no-replay baseline, indicating that grid-pattern stability might be maintained by external replay input rather than self-sustained within the EC; grid-axis replay produced more robust grid patterns than motion-axis replay at larger grid scales, supporting the view that hippocampal replay directed along the grid axes promotes robust grid-pattern formation.

### Two-fold periodicity in trajectory deviation aligns with EC grid orientation

Beyond the neural and modelling results above, we next questioned whether the replay and grid-like activity was also reflected in participants’ navigation behaviour. Specifically, for each connection, we quantified the instantaneous movement speed of the cursor (deg/ms) and the trajectory deviation, defined as the perpendicular distance of the cursor from the veridical direction, as two trajectory measures. To ensure each trajectory contained the complete movement points during connection, movement trajectories were normalized to a 0–1 scale (percentage of trajectory progress).

Both measures rose and fell smoothly across the connection, and their peaks were temporally offset, with speed peaking earlier than deviation (Fig. 5a; 39.7% vs 50.7% of trajectory progress; t(24) = 7.09, p = 2.5 × 10⁻⁷, Cohen’s d = 1.42). This early rise in speed followed by a later peak in deviation suggests that participants first accelerated and then deviated from the veridical direction, and these directional adjustments might be structured by the underlying grid geometry. To examine this hypothesis, the speed and deviation were computed at each percentage point along the trajectory across 120 motion directions, and the FFT power from one- to eight-fold periodicities was extracted.

**Figure 5.**
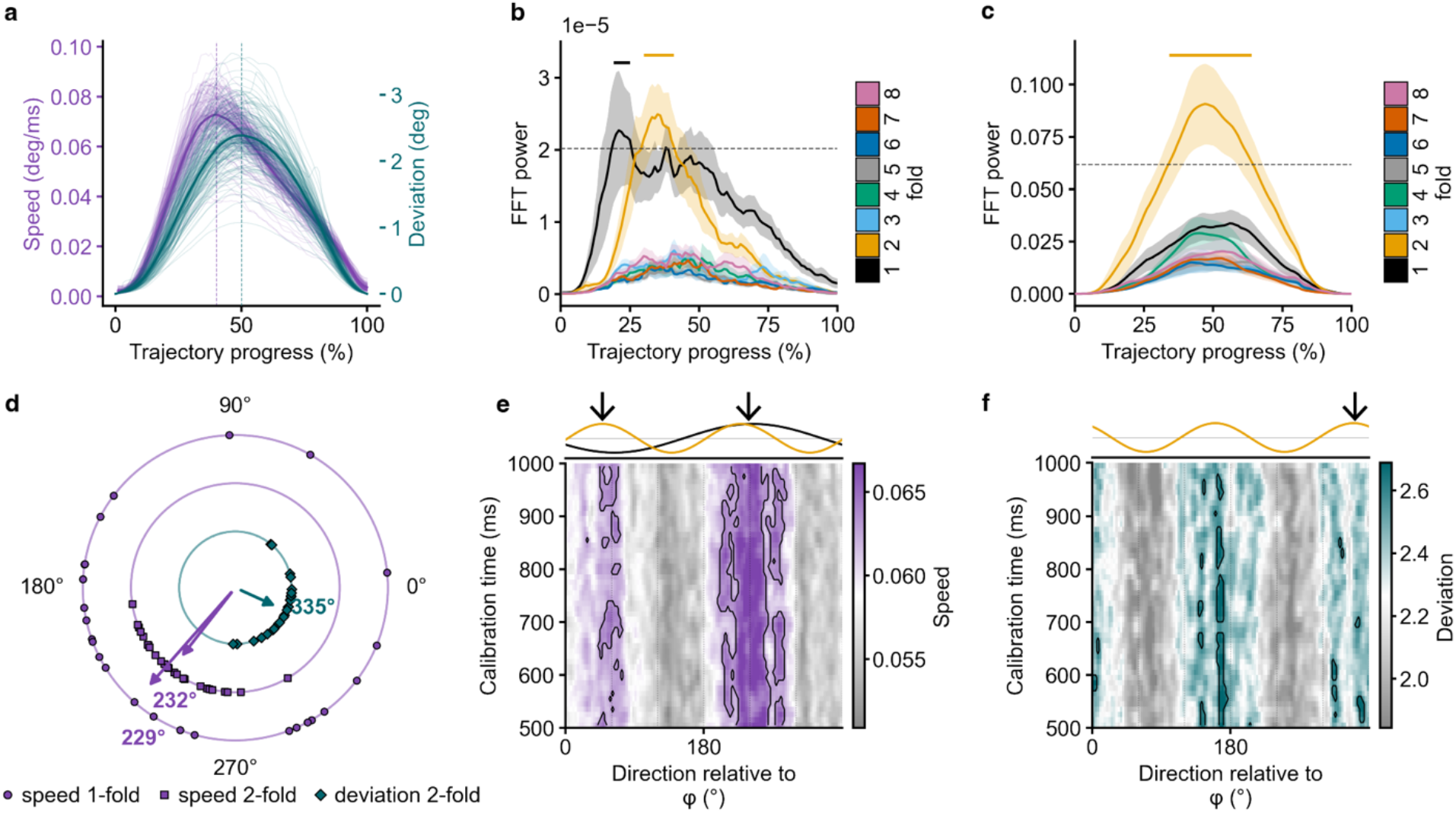
Trajectory deviation selectively aligns with entorhinal grid axes during connection. (a) Speed (purple) and trajectory deviation (teal) profiles along the normalized trajectory of each pairwise connection. Thin lines, per-direction profiles averaged across participants; thick lines, across-participant mean ± s.e.m. Speed peaked earlier in the trajectory than deviation (peak progress 39.7% vs 50.7%; per-subject paired t-test, t(24) = 7.09, p = 2.5 × 10⁻⁷, Cohen’s d = 1.42). Note: the direction sequences of both speed and deviation were realigned to each participant’s grid orientation φ to enable further across-participant comparison. (b-c) Direction-tuning FFT power of movement speed and deviation as a function of trajectory progress for one- to eight-fold periodicities. Dashed line, permutation-based significance threshold. Speed showed significant one-fold (19–25%) and two-fold (29–41%) power early in the trajectory, whereas deviation showed significant two-fold power later (35–63%). (d) Per-participant phase of the grid-orientation–realigned direction tuning, for speed (one-fold, circles; two-fold, squares) and deviation (two-fold, diamonds). The one- and two-fold speed phases did not differ across participants (per-subject phase difference: V-test for clustering toward 0°, u = 2.41, p = 0.008). Markers, individual participants; arrows, across-participant mean resultant vector. The deviation two-fold phase was significantly aligned with the grid axes (mean = 335°, R = 0.48; V-test for clustering toward 0°, u = 2.23, p = 0.013), whereas neither speed periodicity was aligned with the grid axes (one-fold = 229°, R = 0.44, u = −2.47, p = 0.99; two-fold = 232°, R = 0.63, u = −1.01, p = 0.84). (e-f) Grid-orientation–realigned tuning as a function of motion direction relative to φ (x-axis) and grid-orientation calibration time (y-axis), for speed (e) and deviation (f). Speed modulation peaked near 60° and 240°, whereas deviation modulation peaked near 0°, indicating alignment with the grid axes. Top, fitted periodic schematic with arrows marking the modulation peaks. Contours mark significant clusters (FWE-corrected p < 0.05).

Speed showed significant one-fold and two-fold power (Fig. 5b; one-fold: 19–25% progress; two-fold: 29–41% progress; FWE-corrected, p < 0.05), both confined to the early period during connection, whereas deviation showed significant two-fold power later (Fig. 5c; 35–63%). We refer to the segments of the connection in which FFT power exceeded the permutation threshold as significant-periodicity windows. These windows matched the behavioral dynamics in Fig. 5a in two respects. First, within each measure, the window of significant periodicity covered that measure’s own behavioral peak. Second, across measures, periodicity in speed reached significance earlier along the connection than periodicity in deviation, consistent with the ∼10% lead of the speed peak over the deviation peak (Fig. 5a). Notably, this temporal segregation, speed periodicity in the early window and deviation periodicity in the late window, mirrored the temporal profile of replay activity (Fig. 1g; before 500 ms from stimulus onset) and the emergence of entorhinal six-fold grid coding (Fig. 1l; after 500 ms).

A two-fold modulation implies that, out of the full circle of directions that cover 360°, speed and deviation were each selectively enhanced for one direction and its 180°-opposite, paralleling the axis-opposite (forward/reverse) organisation of replay (Fig. 3). Therefore, we next sought to identify which directions these modulations encoded. In doing so, the direction tuning to each participant’s grid orientation φ was realigned, and the per-subject phase of the speed one- and two-fold modulations and of the deviation two-fold modulation were extracted. For the speed modulations, the across-participant mean phases of the one- and two-fold modulations were closely matched (one-fold = 229°, two-fold = 232°), and the per-subject one-fold–two-fold phase difference clustered significantly around zero (Fig. 5d; V-test for clustering toward 0°, u = 2.41, p = 0.008), indicating that the one- and two-fold phases were aligned with each other, and neither was aligned to the grid axes (V-test for clustering toward 0°: one-fold u = −2.47, p = 0.99; two-fold u = −1.01, p = 0.84). By contrast, the deviation two-fold phase clustered at the grid axis (335°; Fig. 5d, teal arrow, V-test toward 0°: u = 2.23, p = 0.013), indicating that deviation was selectively enhanced along participants’ grid axes.

To further validate the robustness of speed and deviation periodicity and examine how their directional phase depended on the grid-orientation estimate, we recomputed the realigned speed and deviation across directions using the grid orientation extracted at each 500–1000 ms time point. Given that the speed one- and two-fold phases were not significantly different (Fig. 5d), their fluctuation maps were averaged. Both resulting maps showed significant directional clustering (Fig. 5e & f; FWE-corrected p < 0.05). Speed formed significant clusters around 60° and 240° (Fig.5e), whereas deviation formed significant clusters near 0° and 180° (Fig.5f), that is, centred on the reference grid axis φ. Both panels exhibited vertical bands, indicating that the directional selectivity of speed and deviation remained stable across the grid-orientation estimate.

These results show that the hexagonal grid-axis organisation observed in replay and EC activity is also expressed in human navigation behaviour. During each connection, participants first accelerated, producing an early, non-grid-aligned speed modulation; they then deviated strongly from the second half of the time window onward, and these adjusted heading directions became aligned to the grid orientation φ. This early-speed-then-late-deviation sequence, together with the selective alignment of deviation to the grid axis, parallels the replay-to-grid temporal pattern documented in the neural data (Figs 1–3).

## Discussion

The present study characterized hippocampal replay and entorhinal six-fold grid coding within a single human MEG task that spanned the full directional space, and found that replay is geometrically organized along the grid axes, with forward and reverse replay unfolding along the two opposite directions of each grid axis. In a continuous-attractor network model, replay directed along the grid axes produced more robust grid patterns, with significantly higher gridness, than replay along the movement axis or without replay, indicating that grid-axis replay supports stable grid formation. These findings were further supported by navigation behaviour. Both movement speed and trajectory deviation exhibited two-fold directional periodicity, with their two preferred directions separated by 180°, and the phase of deviation aligned with the grid orientation, indicating that the grid orientation became aligned with the goal location toward the end of each connection. These results support the view that replay and grid coding are coordinated in geometry.

Our central directional finding is that forward and reverse replay unfold along the two opposite directions of the same grid axes. This extends previous reports of temporal coordination between HPC and EC replay to the directional domain ^15,16^ and of offline hippocampal ripples predicting later grid-like coding in humans^19^. This organization points to an operation required for self-centred navigation. The HPC encodes self-location through place cells^30^, the EC provides a global metric through grid cells^10,31^, and flexible behaviour requires binding ’where I am’ to this global metric^32,33^. Reverse replay (looking back toward the origin) and forward replay (looking ahead toward the destination), engaged one after the other along the same metric axis, may be the neural expression of this binding of self-location to the global metric.

Importantly, this 180° opposition does not imply that reverse and forward replay selectively encode the 0–180° and 180–360° ranges, respectively, as the directional binning in Fig. 3a might suggest, for three reasons. First, the assignment of forward and reverse replay to opposite halves was obtained after aligning to each participant’s grid orientation, a reference that is blind to the actual spatial direction during the connection. Second, the two replay types were characterized at temporally separate windows, with reverse replay earlier (around 150 ms) and forward replay later (around 500 ms); the 180° opposition therefore reflects a relationship between the two replay types across these windows rather than a continuous relationship maintained throughout the connection. Third, the absolute phases of forward and reverse replay were distributed near-uniformly across 0–360° (Fig. 3f), and only their difference was reliably clustered near 180° across participants. Our analyses therefore do not support an absolute directional representation for either forward or reverse replay, and only the 180° opposition between the two is interpretable.

Our results indicate a pathway from the HPC to the EC, in which hippocampal replay activity participates in organizing entorhinal grid coding. This is consistent with the dependence of grid firing on excitatory hippocampal drive^14^ and with the HPC leading the EC by approximately 11 ms during replay^15^. Classical and oscillatory-interference models, by contrast, emphasize the opposite direction, in which entorhinal grids provide metric input to hippocampal place cells^34^. These observations suggest that the EC and HPC form a recurrent loop with bidirectional information flow. This loop also clarifies an apparent temporal mismatch. Although six-fold grid coding becomes significant in the EC only at 500–1000 ms, the directional organization of replay is already present earlier, before 500 ms. We propose that the grid orientation is not generated at 500 ms and then read out by replay; rather, the early, 180°-opposed forward and reverse replay may establish the grid orientation, so that the grid axes come to align with the replay directions, and the six-fold firing at 500–1000 ms is the consolidation and read-out of this orientation in the EC. Replay and grid are thus not related by sequential read-out but are mutually constructed within the loop, which also accommodates the goal-dependence of grid orientation and the simultaneous engagement of all three axes by replay. Our continuous-attractor network provides an instance of this loop, in which axis-aligned, place-driven input within a recurrent network establishes and stabilizes the six-fold read-out (Fig. 4). Our evidence for a replay-to-grid influence rests on the temporal precedence and directional alignment of replay relative to the entorhinal six-fold code, together with a network model and convergent behaviour; this evidence remains correlational, and establishing causality will require directly manipulating replay, for example by disrupting hippocampal sharp-wave ripples during navigation.

The grid pattern, with its three axes, intrinsically encodes orientation rather than a directed vector, because the six-fold code is 60°-periodic and each axis is 180°-symmetric. To specify a direction, this axial code must be combined with a directional signal, and the EC already carries such signals, with conjunctive grid × head-direction cells conferring a heading on the grid axes^23^, and vector coding being a predominant form of position coding^35^. We propose that forward and reverse replay, by breaking the 180° symmetry of each axis and assigning values to its two opposite directions, convert the undirected axial metric into a directed vector; anchored to an allocentric grid with the self as origin, this constitutes a self-to-goal mapping vector, in which reverse replay corresponds to the backward propagation of experience and value and forward replay to prospective planning^1,2^. This framework also offers a new account of spatial reorientation. After being disoriented, for example by passive rotation, rats in rectangular enclosures^36^ and human toddlers^37^ reorient preferentially by the global geometry of the enclosure rather than by salient local features, and systematically confuse a target location with its 180°-rotational equivalent. We suggest that this behaviour corroborates the present picture in two respects. First, after disorientation the brain represents global orientation, the axial geometry, before local detail, indicating that an orientation representation already exists in the brain. Second, this orientation is not converted into a directed vector that would resolve the 180° ambiguity, possibly because replay has not yet integrated local feature information into the metric coded by entorhinal grid cells; consistent with this, disrupting hippocampal sharp-wave ripples impairs spatial memory and learning^38,39^. The reorientation behaviour thus corresponds to a state in which orientation is present but the replay-conferred vector is absent, whereas in our task forward and reverse replay accomplish this conversion from an axial orientation to a directed vector, yielding a unique heading.

The present results suggest that replay does not only support memory and planning but also contributes to organizing the spatial metric itself. Together with the findings that grid-like activity codes non-spatial two-dimensional metrics such as conceptual and olfactory spaces^11,12^, the organizing role of hippocampal replay over grid geometry may generalize as a mechanism for constructing abstract cognitive maps and supporting flexible navigation.

## Methods

### Participants

Twenty-five right-handed university students with normal or corrected-to-normal vision were recruited from Tsinghua University (mean age = 25.2 [SD = 3.78], 18 male, 7 female). All participants had no history of psychiatric or neurological disorders and provided written informed consent prior to testing. The study protocol was approved by the Research Ethics Committee of the Faculty of Psychology, Beijing Normal University (approval number: 202003180020). Data were collected while the laboratory was at Beijing Normal University. The sample size (n = 25) is in line with prior human MEG studies of hippocampal replay^4,5^, in which sequenceness has been reliably detected at comparable sample size.

### Experimental procedure

Visual Stimuli. Four nonmeaningful stimuli were used as visual stimuli. They were manually selected from the original Omniglot handwritten-stimulus dataset^40^ following two criteria: (i) each stimulus was easy to recognize (e.g., few strokes), and (ii) stimuli within a set were clearly distinguishable from each other. After this curation, 72 candidate stimuli were retained, from which a unique set of four was assigned to each participant for each session. Character positions were determined by one of 120 evenly spaced directions. Stimuli were placed on a virtual spherical surface, with a Unity perspective camera (participants’ viewpoint; field of view = 70°) at the centre, and each stimulus oriented toward the camera at a fixed distance. Stimuli were rendered at 105 × 105 px on the in-bore 1024 × 768 MEG projection screen.

### Paradigm

Participants performed a relational-memory paradigm: they learned the pairwise spatial sequence between four visual stimuli arranged in a two-dimensional latent space, with the sequence introduced through an instruction cue at the start of each session. The space is termed ’latent’ in the sense that, with the camera at the centre of the spherical stimulus surface, only a portion of the stimuli was visible in any one frame; the full pairwise map was built by integrating these partial views across mouse-driven rotations. Each participant completed five consecutive MEG recording sessions, each consisting of 120 four-stimulus sequence trials. On each MEG trial, participants viewed a starting stimulus and were asked to make the connections A→B, B→C and C→D as accurately and as fast as possible, after which corrective feedback was displayed. Trial-onset and choice events were time-locked to the MEG recording. This paradigm gives 5 × 120 × 3 = 1,800 connection events for each participant in total. Behavioral responses were collected through a fibreoptic MEG-compatible mouse (NAtA Technologies, fMRI Mouse System, model FOM-2B-10B) with two standard buttons (Left and Right); only the left button was used for the connection click.

### Data Acquisition

MEG acquisition. Neuromagnetic signals were continuously recorded with a 275-channel whole-head axial gradiometer system (DSQ-3500, CTF MEG, Canada) at a sampling rate of 600 Hz. A third-order synthetic gradiometer and linear drift correction were applied to remove environmental noise. Head position within the MEG helmet was monitored using three head-position indicator (HPI) coils attached to the nasion and bilateral preauricular points, enabling co-registration with the sensor coordinate system. To minimize head movement, customized MEG-compatible chin-rest equipment was used (Fig. S6; Written informed consent for image publication was obtained), ensuring that movement did not exceed 2 mm during recording.

### Anatomical MRI data acquisition

After the MEG sessions, each participant underwent anatomical MRI scanning on a 3T Siemens Prisma scanner (voxel size: 1 mm³; flip angle: 7°; TE: 3.37 ms; TR: 2200 ms; field of view: 224 × 256 × 192 mm³). During the MRI scan, the same three magnetic coils worn during MEG recording were marked with MRI-visible fiducials, allowing spatial registration between source-space anatomy and MEG sensor positions.

### MEG Preprocessing

MEG data were preprocessed using the MNE-Python toolbox^41,42^. For each participant, raw CTF data from the five MEG sessions were high-pass filtered above 0.5 Hz using a FIR firwin filter, and were further downsampled to 100 Hz. To verify data quality and remove possible contamination from artifacts (cardiac activity, eye movements, blinks, environmental noise), an independent component analysis (ICA) was performed. Each participant’s T1-weighted MRI was processed with FreeSurfer recon-all^43,44^ to obtain cortical surface reconstructions and skull-stripped brain volumes. Inner-skull surfaces were extracted with mne.bem.make_watershed_bem and used to build a single-compartment boundary element model (ico = 4, conductivity = 0.3 S/m). MEG–MRI coregistration was performed in MNE-Python by aligning the three magnetic coil locations and the digitised head-shape points to each participant’s anatomical surface. Volumetric source spaces were created with 5-mm isotropic spacing within the inner-skull surface, and surface source spaces with ico5 spacing (∼4.9 mm). Individual source estimates were morphed to the MNI template for group-level statistics.

### Replay analysis

Four binary one-vs-rest L1-regularised logistic-regression decoders (sklearn LogisticRegression, C = 0.005, fitted independently at each 10 ms time-step from 0 to 1000 ms post-stimulus) were trained on MEG sensor data to discriminate the four stimuli. Each trial’s features were standardised against a 50-ms pre-event baseline. For the subsequent replay analyses, the peak-performance decoder for each symbol was extracted at 300 ms post-stimulus.

Decoder performance was estimated by 5-fold cross-validation. The trained peak-performance decoder for each symbol was then applied to MEG activity acquired during each connection (0–1000 ms relative to connection onset), yielding a stimulus-specific reactivation probability time-course.

Sequenceness, the directional structure of stimulus reactivation across time, was quantified using temporally delayed linear modelling^4^. For each connection, the four reactivation time-courses were entered into a regression model at successive time lags (10–100 ms in 10 ms steps), separately for the forward and reverse transition matrices. Source-space sequenceness within the hippocampus was estimated at three time points of interest (150, 300, 500 ms) using the source-localization procedure detailed below. To test whether replay events were anchored to grid-cell axes, TDLM was re-performed for each direction (i.e. trial), and the resulting replay activity was projected onto twelve evenly-spaced movement directions and binned into “aligned” (within ±15° of one of the six 60°-spaced grid axes) and “misaligned” direction relative to the grid orientation extracted from the concurrent entorhinal source-space activity.

### Sinusoid analysis

Following the established grid-cell analysis approach in human MEG and fMRI^11,24^, we tested for a population-level six-fold modulation of EC activity as a function of direction θ using the cosine modulator M(θ; φ) = cos(6 (θ − φ)), where φ is the participant’s preferred grid orientation. EC was manually delineated on a 2mm T1 MNI template using established protocols^45,46,47,48^ and the delineating software ITK-SNAP (www.itksnap.org). φ was estimated from EC source activity in a five-fold cross-validation procedure: the trials of each session were partitioned into five folds; in each cross-validation iteration, four folds were used to estimate φ (by fitting cos(6 θ) and sin(6 θ) regressors at every voxel × time-point), and the held-out fold was used to evaluate the cosine-aligned EC activity at the estimated φ. The held-out cosine-aligned EC activity was then averaged across voxels, sessions and folds for each participant, yielding an individual-participant time-dependent six-fold modulation amplitude. For specificity, the same procedure was repeated with control fold-N modulators (N ∈ {4, 5, 7, 8}) to confirm that the modulation was selective to the six-fold periodicity. Source-space significance for the bilateral EC ROI was assessed analogously with cluster-based permutation in source space (initial threshold p < 0.05, cluster permutation p < 0.05, two-tailed).

### FFT analysis of replay direction tuning

To characterize the spectral structure of replay tuning along directions, we computed FFT power of replay activity across 120 directions after grid-orientation realignment. For each participant, symbol, and cross-validation fold, the realigned forward and reverse 120-direction signals were combined and resampled to N_d_ equally-spaced direction bins. To balance bin-by-bin signal-to-noise (smaller N_d_ averages over more original direction samples for each bin) against frequency resolution, N_d_ was chosen as the smallest integer satisfying N_d_ / 2 > 6, yielding N_d_ = 14 (n_half_ = 7), so that folds 0–6 are all resolvable. After resampling, the signal was linearly detrended along the direction axis and tapered with a Hanning window prior to a discrete Fourier transform; FFT power |FFT|² was retained for folds 0–6. Power was first averaged across cross-validation folds and symbols for each participant, then across participants. To test for direction-asymmetric vs. axis-symmetric components, two complementary analyses were applied at the 120-direction level:

i. Forward and reverse replay analyzed independently. Each replay type’s 120-direction signal was analyzed through the FFT pipeline separately. This analysis is sensitive to direction-asymmetric components (fold-1, fold-3).
ii. Forward and reverse replay averaged. The two replay-type signals were averaged at the 120-direction level (combine_fn(f, b) = (f + b) / 2) before resampling and FFT. Under this average, the direction-asymmetric components (fold-1 and fold-3) are suppressed by the anti-phase relation between forward and reverse replay, while the axis-symmetric component (fold-6) is preserved by their in-phase relation; this isolates the axis-symmetric six-fold grid signature.

For the FFT phase analysis (Fig. 3f), the same FFT pipeline was applied to forward and reverse replay independently in raw motion-direction space (without grid-orientation realignment and without averaging), yielding an individual-participant complex FFT spectrum at each fold. The angle of the complex FFT coefficient at the fold of interest was used as that subject’s phase. Group-level Rayleigh tests were applied to the individual-participant phase distributions for forward, reverse, and the within-subject reverse−forward phase difference.

### Simulation of multi-scale hexagonal pattern

Continuous-attractor network (CAN) simulations were implemented in PyTorch (Python 3.10). Each CAN comprised n × n = 101 × 101 grid-cell units; recurrent two-dimensional symmetric (W) and asymmetric (W_asy_) weight matrices were generated using the Ψ-function definition of Fuhs and Touretzky^29^ with N_w_ = 3, amplitude = 1/3, and spatial frequency ω = 2π/σ for grid scale σ. The voltage of each grid-cell unit ξ_i_ was initialised to zero and updated at each simulation step by computing the symmetric and asymmetric drives below.

Symmetric drive at unit i, given the index set E(t) of currently excited units (those with ξ > 0):

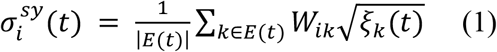

Asymmetric drive averaged across a direction set D, where the shift operator S_d(k)_ shifts the index k by an offset along direction d (with bilinear interpolation to obtain sub-pixel accuracy):

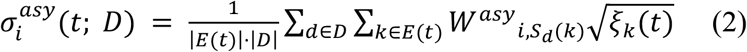

Discrete dynamics update with gain g:

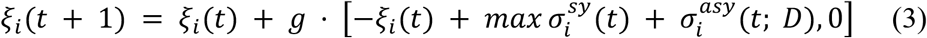

The same update equation applied to all three replay conditions; they differed only in the direction set D and the gain g. During motion phases (in all three conditions), the gain was set to g_motion_ = 0.05; during reverse-replay and forward-replay phases (in the two replay conditions), the gain was set to g_replay_ = 1.0. Direction sets D for each phase are specified in the next paragraph. The 20-fold higher gain during replay (g_replay_ /g_motion_ = 20) implements the time-compressed amplitude amplification observed in empirical replay events.

To probe the grid-scale × replay interaction, eleven grid scales were simulated: σ ∈ {20, 25, 30, 35, 40, 45, 50, 55, 60, 65, 70} pixels (≈ 20%–69% of the representational space). To capture the three hexagonal axes of biological grid cells, twelve evenly-spaced movement directions were used for each simulation (0°, 15°, 30°, …, 165°).

### Simulation of hippocampal replay

Three replay conditions were compared. In the no-replay condition, the CAN was driven by movement-direction input only; the simulation consisted of consecutive blocks of constant-direction movement updates with no additional sequential drive. In the motion-axis replay condition, movement blocks were interleaved with two short place-cell-driven replay phases for each cycle: a reverse-replay phase in which the activated place-cell sequence applied 180° opposite to the current movement direction (θ_motion_ + 180°), followed by a forward-replay phase along the current movement direction (θ_motion_). In the grid-axis replay condition, the reverse and forward replay phases were instead aligned with the three nearest hexagonal axes of the contemporaneous grid orientation: for reverse replay, the three hex directions surrounding (θ_motion_ + 180°) were co-activated; for forward replay, the three hex directions surrounding θ_motion_.

For each simulation we ran 8 cycles of motion → reverse-replay → forward-replay, with each phase within a cycle lasting 10 simulation steps. The number of cycles was chosen so that the total simulation step budget (240 steps) was matched across the three replay conditions (motion-axis and grid-axis replay: 80 motion + 80 reverse-replay + 80 forward-replay steps; no-replay: 240 motion-only steps). The protocol was repeated for each of the 11 grid scales × 12 movement directions, yielding 132 independent simulations for each condition (396 simulations in total).

### Grid score computation

The spatial regularity of hexagonal patterns, as indexed by grid score, was computed with the following procedures introduced previously^23,50^. The spatial autocorrelation of each hexagonal pattern was estimated by computing Pearson’s product-moment correlation coefficient for each representational location (x, y) of hexagonal pattern based on the neural activity *λ*(*x*, *y*) and the spatial lags *τ_x_* and *τ_y_* over *n* locations. The autocorrelogram was further rotated by 30°, 60°, 90°, 120°, and 150°, the similarity between the rotated autocorrelogram and the original autocorrelogram was assessed by Pearson correlation. Grid score was finally derived by subtracting the highest similarity of 30°, 90° and 150° from the minimum similarity of 60° and 120°.

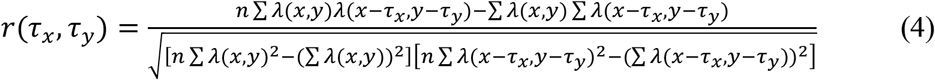

### Statistics

Cluster-based permutation tests were used to assess cluster-level significance. For each analysis, a cluster-forming threshold was applied to the group t-map, contiguous suprathreshold points were combined into clusters, and condition labels were randomly permuted; on each permutation the maximum cluster statistic across the map was retained, and cluster-level significance was defined as the proportion of permutations whose maximum cluster statistic exceeded the observed value, at a corrected threshold of p < 0.05^51,52^. Cluster-forming thresholds and directionality were as follows: p < 0.05 (one-tailed) for the sensor-level replay time-courses, the entorhinal six-fold grid time-course and its control folds, the two-dimensional replay-time × grid-orientation-time map, and the replay FFT power analyses; p < 0.001 (two-tailed) for the whole-brain and entorhinal source-localization analyses; and p < 0.001 for the behavioral direction-tuning analyses. Cluster-level significance was defined as a corrected p < 0.05.

To quantify the significance of hexagonal pattern regularity, the grid score distribution was created using field-shuffling permutation^50,53^, following the same permutation logic described above. The original spatial structures of each hexagonal pattern were shuffled by randomly changing the positions of receptive fields with the local structure (the shape of receptive fields) preserved.

The preferred grid orientation φ (defined modulo 60°) was estimated at every EC voxel and time point. Because grid orientation is 60°-periodic, all circular statistics were computed on 6φ, which maps the 0–60° orientation onto the full 0–360° circle; results are reported and displayed in φ (i.e., divided by 6). For each participant, we tested whether the distribution of grid orientations across EC voxels and time points departed from uniformity with a Rayleigh test (per-participant distributions in Fig. S5), and summarised the strength of clustering by the mean resultant length R. To assess consistency at the group level, each participant’s orientation distribution was first calibrated by subtracting that participant’s own mean orientation; the calibrated distributions were then pooled across participants and again tested for non-uniformity with a Rayleigh test (Fig. 1k).

For the behavioral analyses, the shift in peak trajectory position between speed and deviation tuning was tested with a paired t-test across participants. The one- and two-fold speed phase difference was assessed with the Rayleigh V-test for clustering toward 0°.

## Data availability

The MEG dataset and behavioral data are available from the Science Data Bank at https://doi.org/10.57760/sciencedb.30678.

## Code availability

The task was programmed using Unity (version 6000.0.62f1). Analyses were performed using custom scripts written in Python (version 3.10), with neuroimaging analyses carried out using MNE-Python (version 1.11) and FreeSurfer (version 7.3.2; https://surfer.nmr.mgh.harvard.edu). The task code and analysis source code are publicly available at https://github.com/ZHANGneuro/3D_relational_memory_task.

## Acknowledgements

We thank Fan Wang and Sijia Yang for technical support with the MEG system, and Hang Zhang for providing the MEG-compatible computer mouse used in the experiment. We also thank the MEG Center of the Institute of Biophysics, Chinese Academy of Sciences, for assistance with data acquisition. The authors used Claude to improve the language and clarity of the manuscript. After using this tool, the authors reviewed and edited the content as needed and take full responsibility for the content of the published article.

## Funding

B.Z. discloses support for the research of this work from the China Postdoctoral Science Foundation (2022M710470). J.L. discloses support for the research and publication of this work from the National Natural Science Foundation of China (T2488101).

## Author contributions

B.Z. and J.L. conceived and designed the study. B.Z. and S.T. performed the experiments and collected the data. B.Z. analysed the data and performed the computational modelling. B.Z. and J.L. interpreted the results and wrote the manuscript. J.L. supervised the project. All authors discussed the results and approved the final manuscript.

## Competing interests

The authors declare no competing interests.

---

**Figure S1.**
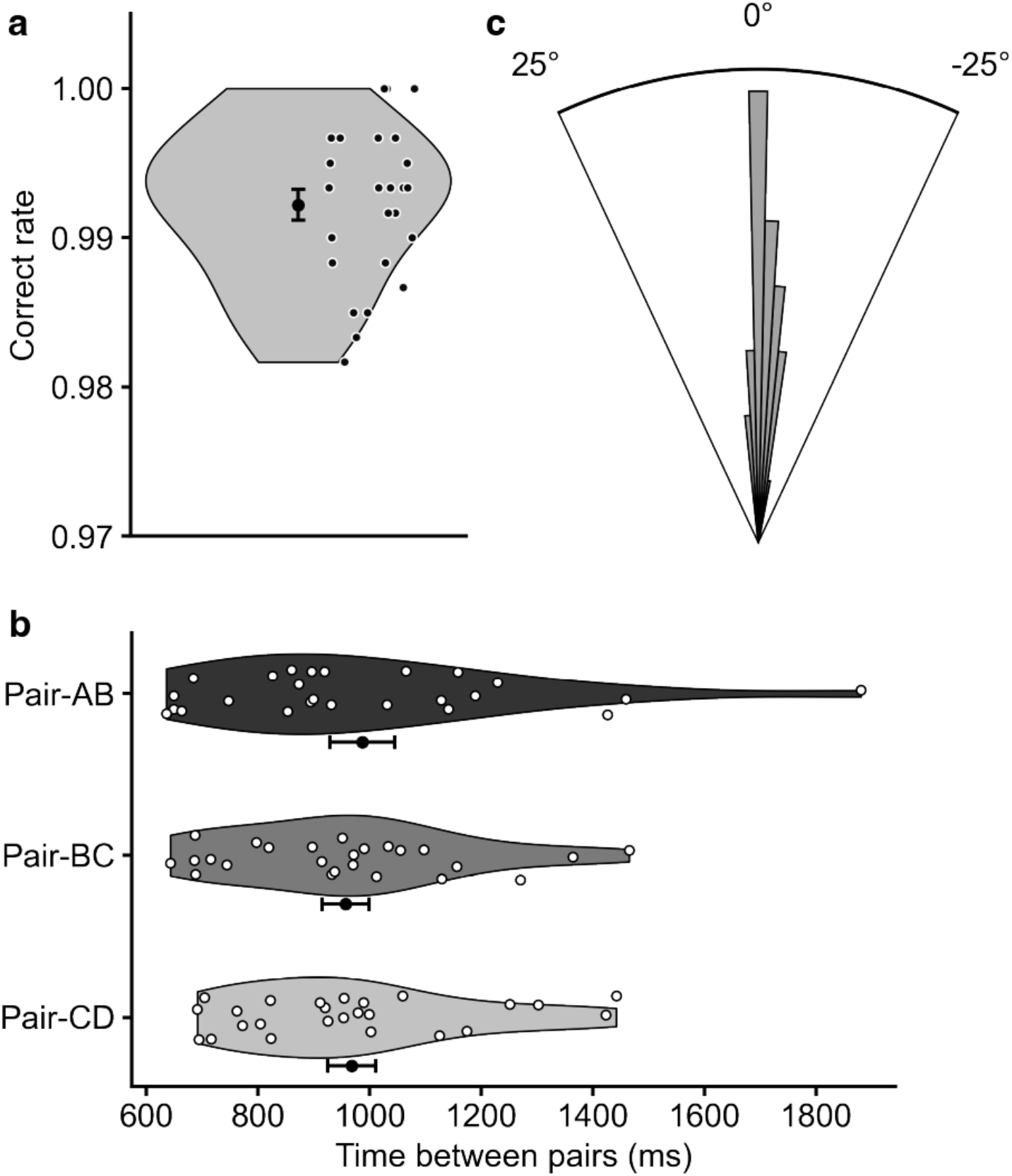
Behavioural performance during connection. (a) Task accuracy across participants; each dot denotes one participant (mean ± s.e.m. = 99.22% ± 0.10%). (b) Connection time duration per stimulus pair (Pair-AB, Pair-BC, Pair-CD). No significant difference among the three pairs (one-way ANOVA, F(2, 48) = 0.46, p = 0.63). (c) Direction errors — distribution of angular differences between participants’ actual movement direction and the instructed direction. The sharp peak near 0° confirms expected task accuracy. ──────────

**Figure S2.**
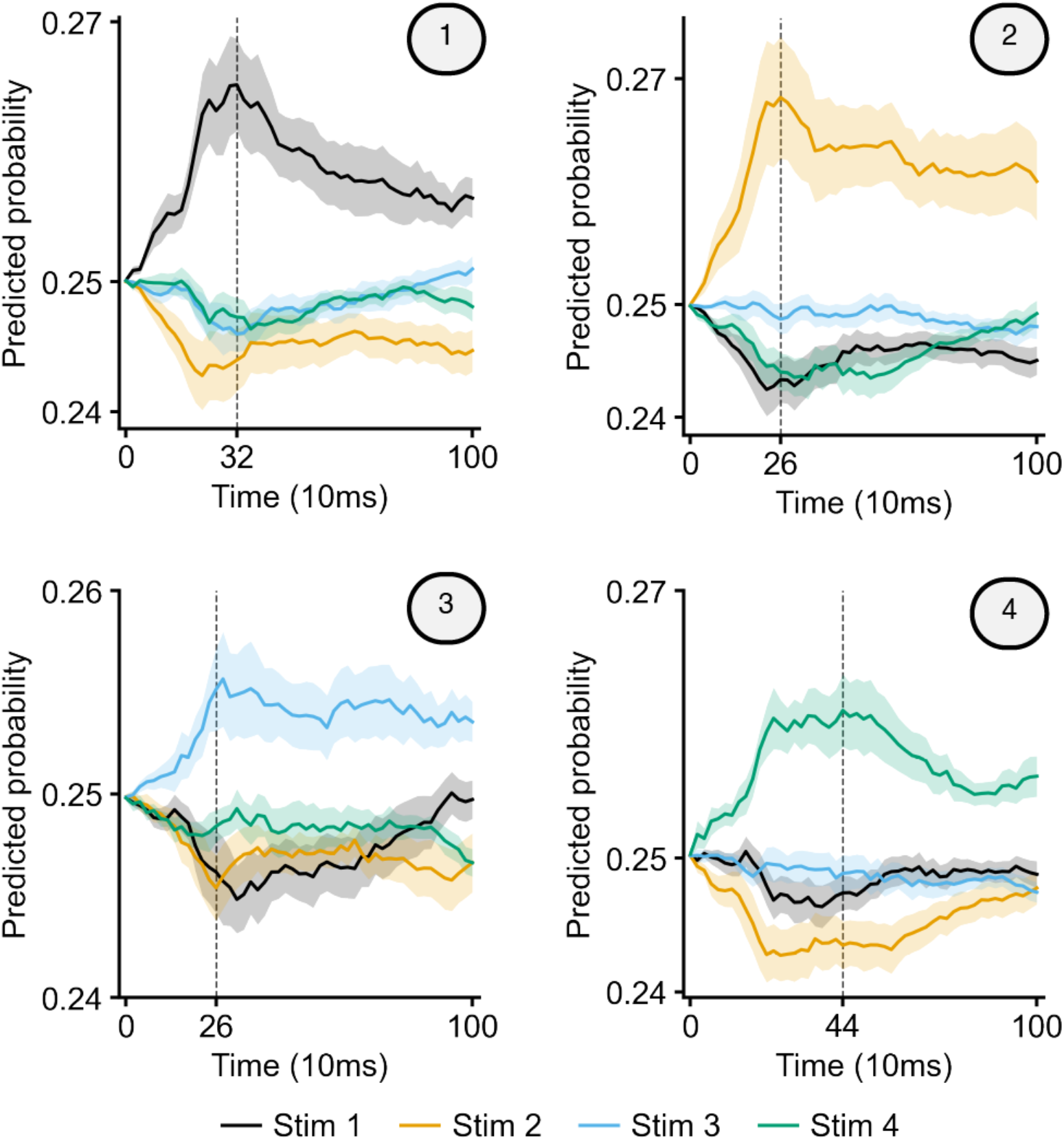
Decoder performance on Omniglot stimulus classification. Each panel shows the predicted probability across time assigned by each of the four classifiers. Shaded band, s.e.m. across participants. In each panel, a vertical line marks the peak timepoint at which the correct class significantly exceeded the mean of the other three classes (paired one-tailed t-test, Bonferroni-corrected across timepoints, p < 0.05). Averaged across the four classifier–class diagonals, decoder output peaked at ∼300 ms post-stimulus (mean predicted probability = 0.262; chance = 0.25).────

**Figure S3.**
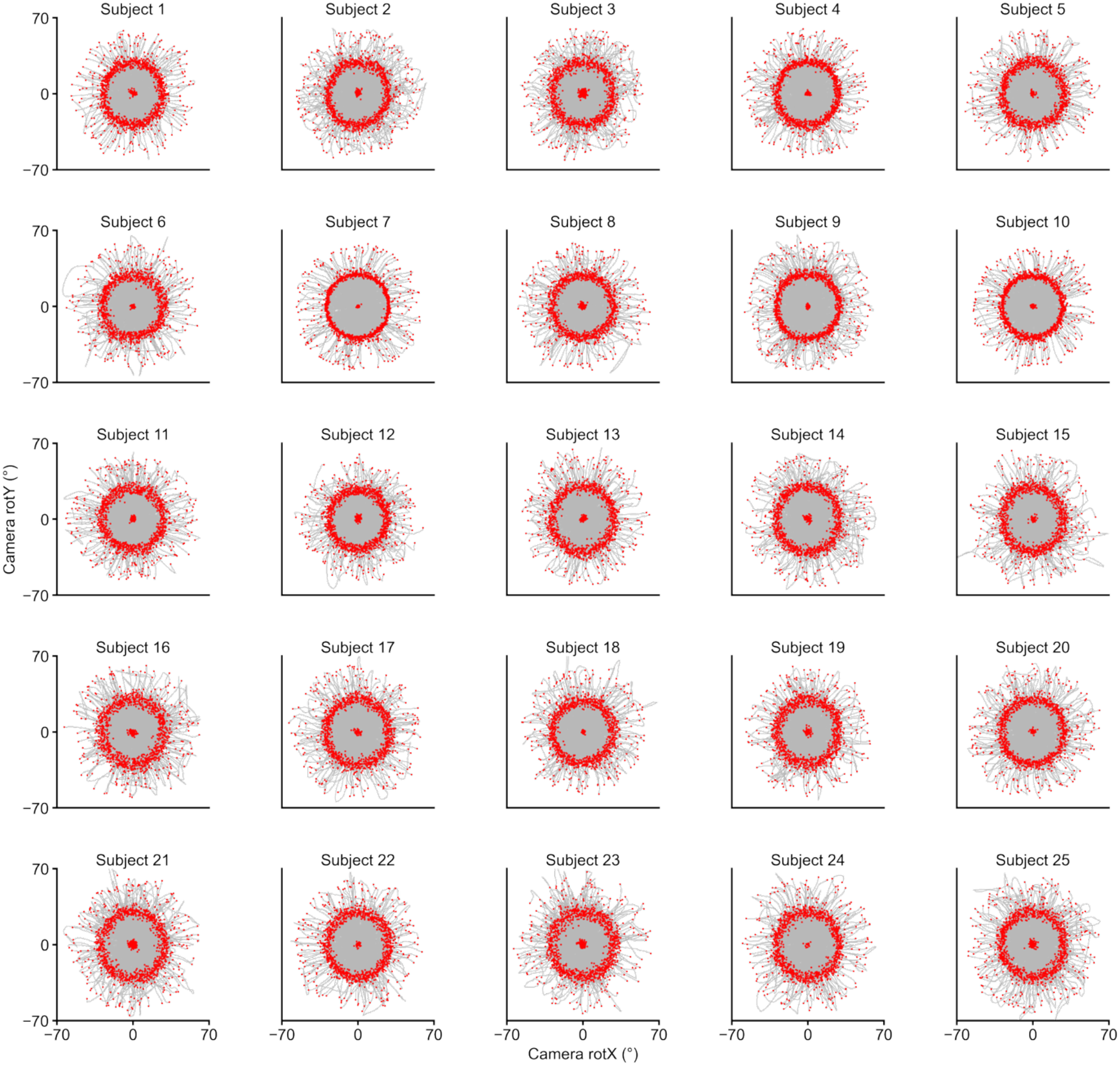
Camera-rotation distribution for each participant during connection. Each panel shows the joint distribution of horizontal (rotX) and vertical (rotY) camera-rotation angles across the entire MEG session for one participant. Gray lines, actual connection trajectories; red dots, samples acquired during the choice phase. ──────────

**Figure S4.**
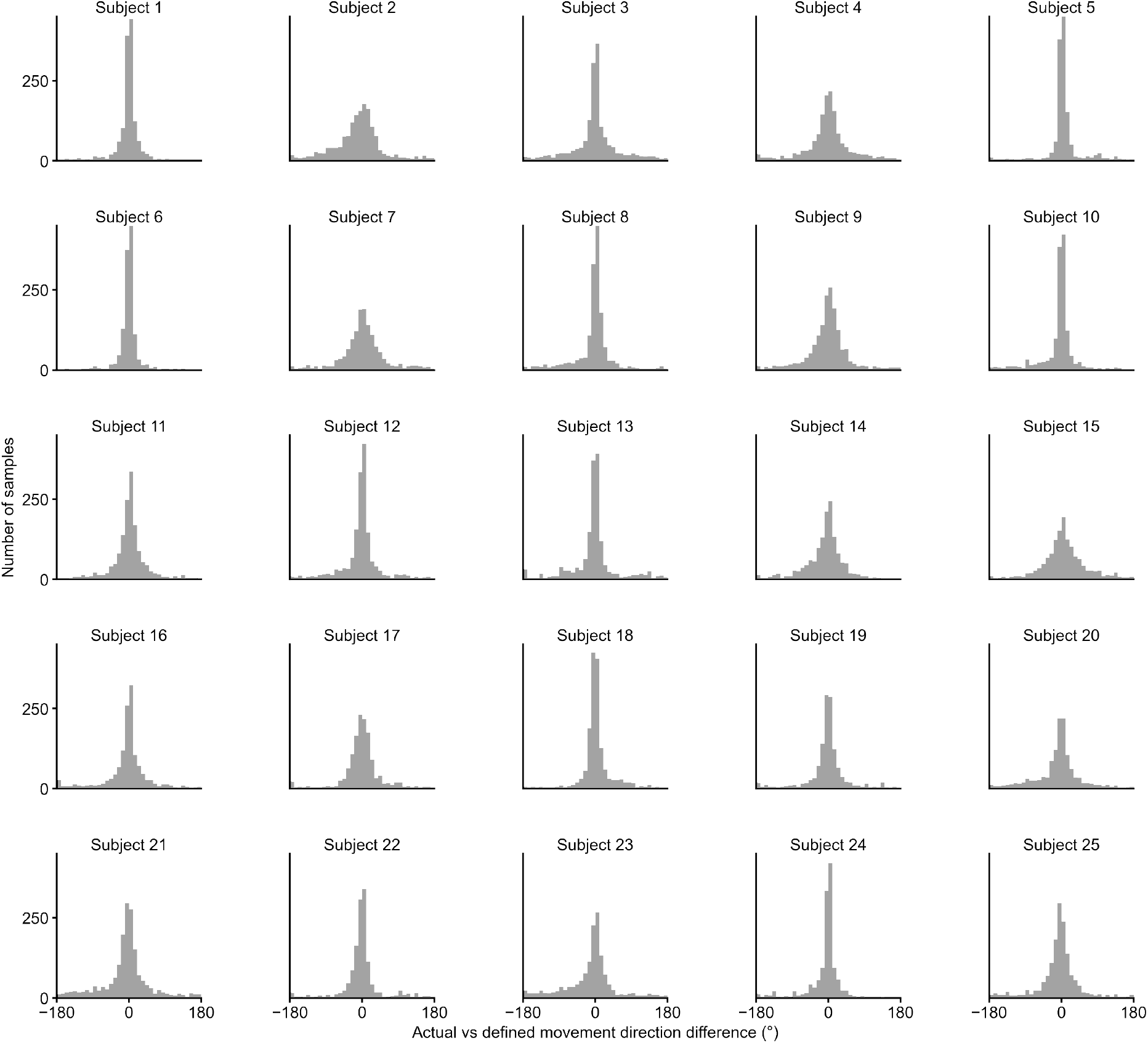
Connection-direction accuracy for each participant. Histogram of the angular difference between each participant’s actual movement direction (from real-time camera rotation logs) and the instructed movement direction.──────────

**Figure S5.**
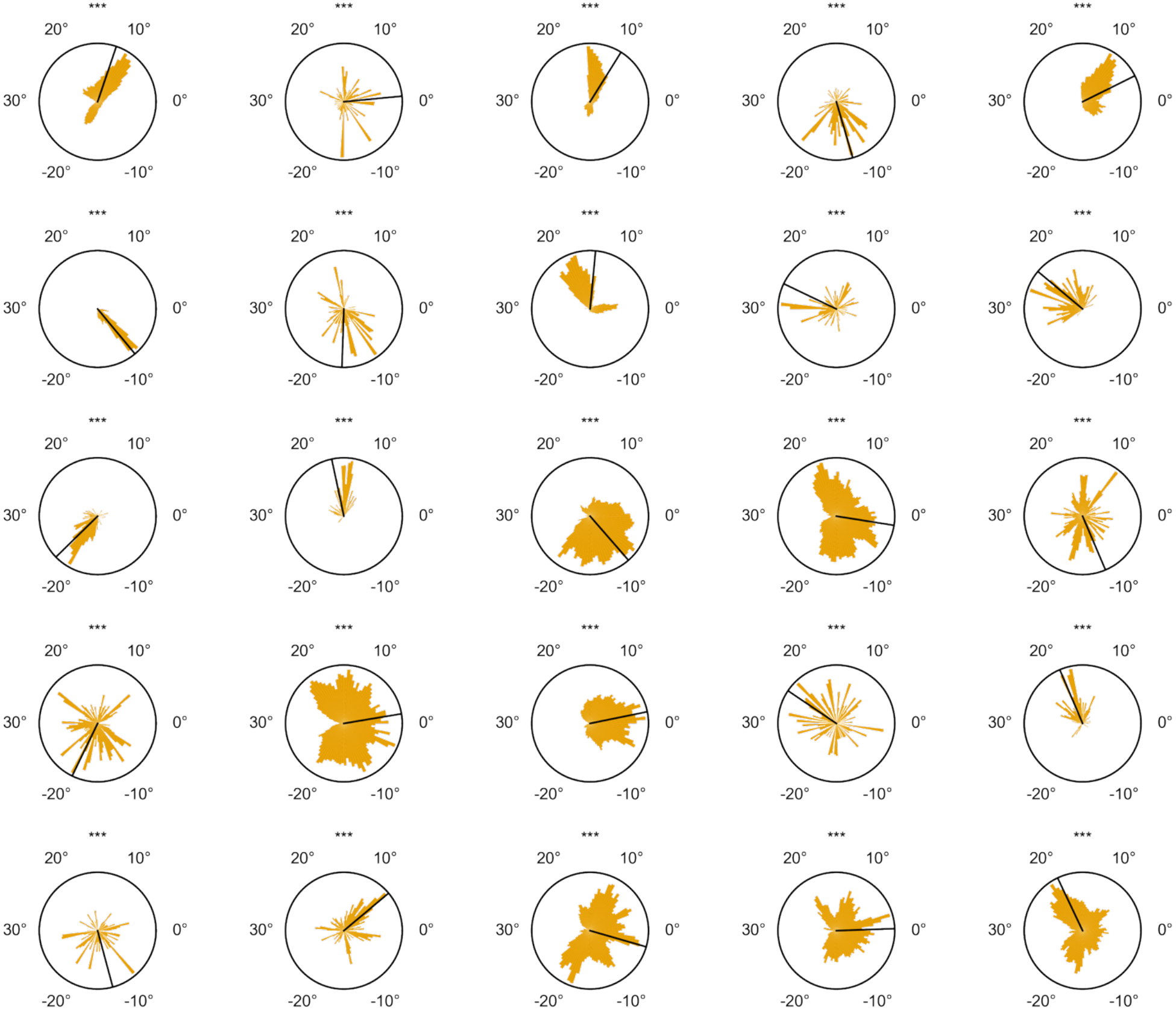
Per-participant grid orientation φ distribution; per-participant Rayleigh test against the uniform null (see Methods).──────────

**Table S1.**
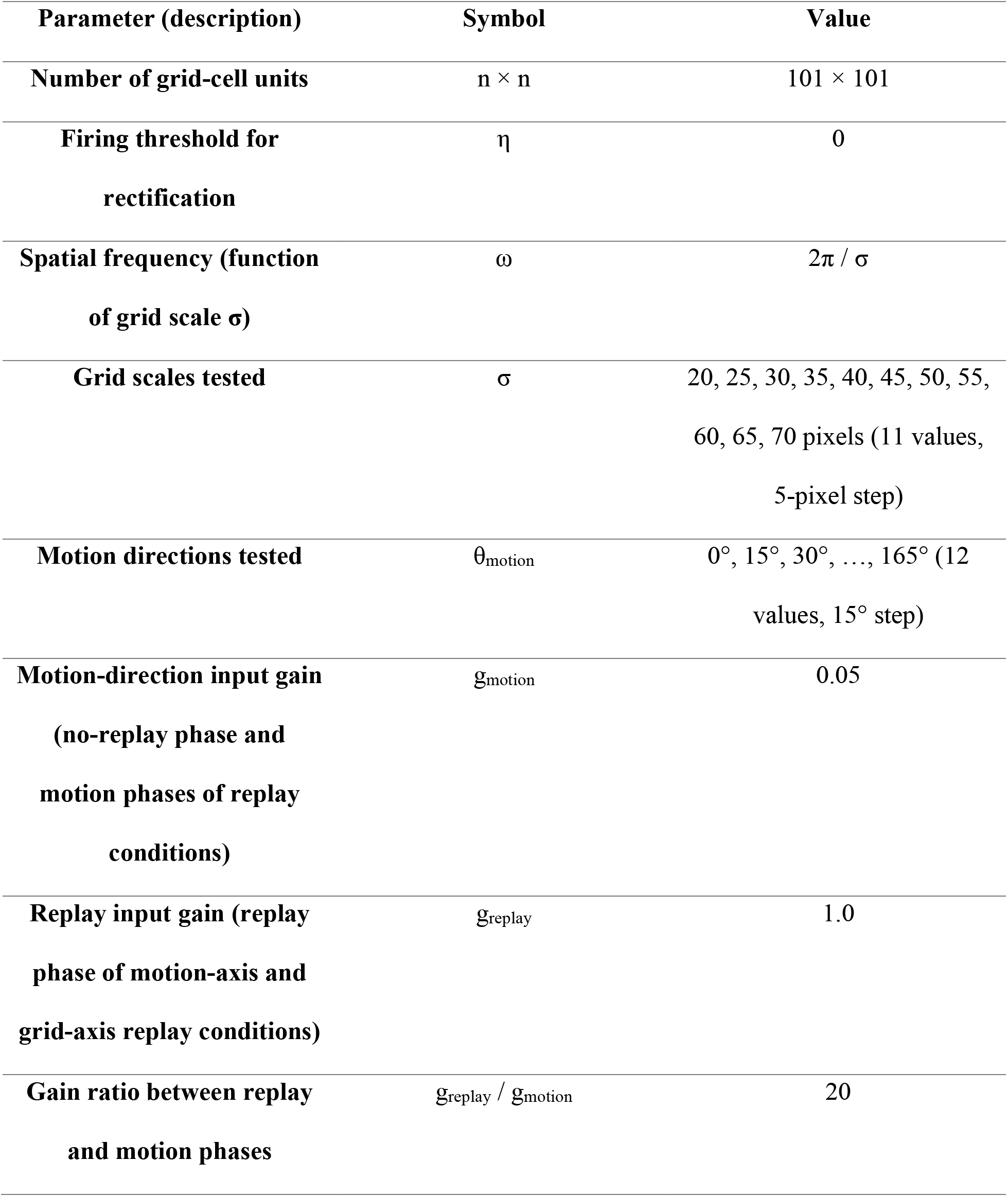
Model parameters.

| Parameter (description) | Symbol | Value |
| --- | --- | --- |
| Number of grid-cell units | $n \times n$ | $101 \times 101$ |
| Firing threshold for rectification | $\eta$ | 0 |
| Spatial frequency (function of grid scale $\sigma$) | $\omega$ | $2\pi / \sigma$ |
| Grid scales tested | $\sigma$ | 20, 25, 30, 35, 40, 45, 50, 55, 60, 65, 70 pixels (11 values, 5-pixel step) |
| Motion directions tested | $\theta_{\text{motion}}$ | $0^\circ, 15^\circ, 30^\circ, \dots, 165^\circ$ (12 values, $15^\circ$ step) |
| Motion-direction input gain (no-replay phase and motion phases of replay conditions) | $g_{\text{motion}}$ | 0.05 |
| Replay input gain (replay phase of motion-axis and grid-axis replay conditions) | $g_{\text{replay}}$ | 1.0 |
| Gain ratio between replay and motion phases | $g_{\text{replay}} / g_{\text{motion}}$ | 20 |

## Notes

### Competing Interest Statement

The authors have declared no competing interest.

### Summary of Updates

Minor revision on abstract, results, discussion, method, and reference.

